# Belowground functional traits predict population temporal stability in grasslands

**DOI:** 10.64898/2026.09.25.754125

**Authors:** Sepideh Golshani, Joana Bergmann, Johanne Gresse, Pierre Liancourt, Maria Májeková

## Abstract

1. Temporal stability of plant populations is a key determinant of species coexistence and ecosystem functioning, yet its trait-based explanation has remained largely aboveground. Whether root functional traits contribute to long-term population stability remains poorly understood, despite roots mediating water and nutrient acquisition.
2. We combined 17 years of vegetation monitoring from 150 temperate grassland plots across three regions in Germany with root functional trait data for 72 common grassland species. Temporal stability was computed as coefficient of variation (CV), with lower CV indicating higher stability. We tested whether root traits predict stability, and specifically which belowground dimensions from resource acquisition/conservation or soil exploration dominate, and whether differences in land use intensity modifies these relationships.
3. Root traits were stronger and more consistent predictors of temporal stability than the classical aboveground traits specific leaf area (SLA) and leaf dry matter content (LDMC). Higher average root diameter (AD), root hair incidence (RHI), specific root length (SRL), and root tissue density (RTD) were generally associated with higher temporal stability, although the strength of these relationships varied among regions.
4. Land use modified selected trait–stability relationships, with the strongest shifts occurring along the mowing gradient. Increasing mowing strengthened the associations of higher RHI and RTD with temporal stability, whereas the associations of AD and root N shifted towards lower stability.
5. Our results identify belowground functional traits as an important missing component of trait-based temporal stability research and advance our understanding of the functional strategies underlying population temporal stability in managed semi-natural grasslands.

## Introduction

Temporal stability of plant populations and communities is of paramount importance for maintaining multiple ecosystem functions over time, such as ecosystem productivity or carbon sequestration (e.g., Lepš et al. 2018; de Bello et al. 2021). Here, temporal stability refers to constancy, which captures the year-to-year fluctuations and describes how invariable an ecosystem property (e.g., abundance) is during a period without particularly extreme perturbations (Doak et al. 1998; de Bello et al. 2021). The temporal stability captures the natural degree of community stability, and as such allows to predict communities’ potential to resist future extreme perturbations (Pimm 1984; de Bello et al. 2021; Liu et al. 2023). Population stability of coexisting species’ is also a key factor from which plant community stability is derived. Understanding the mechanisms underlying temporal stability on different levels of organization is becoming increasingly important under environmental change, which is expected to intensify inter-annual variability in water and nutrient availability across grassland ecosystems worldwide (Knapp et al., 2002; de Vries et al., 2016). At the same time, much less is known about how land-use intensity shapes temporal population stability, particularly whether management modifies the functional trait–stability relationships underlying population dynamics. Alongside the climate-driven environmental variability, land-use management can also alter temporal population stability (Li *et al*., 2023; Zhang *et al*., 2023), yet whether land-use intensity modifies the functional trait–stability relationships underlying population dynamics remains poorly understood.

Over time, species respond differently to environmental conditions depending on their functional strategies. Trait-based approaches can help identify the mechanisms underlying population temporal stability. However, trait-stability relationships have been so far explored mainly using aboveground traits (Wright *et al*., 2004; Májeková *et al*., 2014; Díaz *et al*., 2016; Bruelheide *et al*., 2018; de Bello *et al*., 2021; Conti *et al*., 2023; Gresse *et al*., 2026), particularly traits associated with the resource acquisition-conservation trade-off within the leaf economics spectrum (Wright *et al*., 2004; Díaz *et al*., 2016). In contrast, knowledge about the contribution of belowground traits to long-term temporal stability is virtually non-existing so far despite being consequential as roots are the primary interface through which plants acquire water and nutrients. This gap is particularly important in grasslands, where belowground competition, resource acquisition, and plant-soil interactions strongly influence species responses to environmental variability and consequently species coexistence and diversity in a long term (Mommer *et al*., 2010; Orwin *et al*., 2010; Kroon *et al*., 2012; Laliberté, 2017).

Roots play central roles in water and nutrient acquisition, carbon allocation, and plant-soil interactions, yet root traits have historically been underrepresented in trait-based studies of community dynamics and ecosystem functioning (Laliberté, 2017; Ottaviani *et al*., 2020). Over the last decades, standardized trait protocols, global root trait databases, and improved sampling approaches have substantially expanded the study of root functional ecology across species and allow belowground traits to be linked more directly to plant strategies across species and ecosystems (Bergmann *et al*., 2020; Guerrero-Ramírez *et al*., 2021; Freschet *et al*., 2021; Matthus *et al*., 2025). These recent advances make it possible to ask whether root traits are not only descriptors of belowground form and function, but also predictors of long-term population stability in natural communities.

Different conceptual frameworks have been proposed to describe root functional traits and their ecological consequences. Early approaches often assumed that root traits follow a one-dimensional economic spectrum analogous to the leaf economics spectrum (Reich, 2014; Roumet *et al*., 2016). More recent studies, however, increasingly support multidimensional root trait variation that cannot be fully captured by a single acquisitive-conservative axis (Kramer-Walter *et al*., 2016; Valverde-Barrantes & Blackwood, 2016; Weemstra *et al*., 2016). The root economics space further emphasizes partly independent aspects of root function related to tissue economics and soil resource exploration, including interactions with fungal partners(Bergmann *et al*., 2020; Freschet *et al*., 2021; Matthus *et al*., 2025). Relationships between root traits and ecological performance may also vary strongly across environmental conditions, species pools and timescales rather than reflecting universal belowground strategies (Laughlin *et al*., 2021; Matthus *et al*., 2025).

Despite these advances, it remains unclear whether root functional traits predict long-term population stability in natural plant communities. Most root-trait studies have focused on productivity, resource acquisition or ecosystem functioning over relatively short timescales (Bergmann *et al*., 2020; Freschet, 2021). Long-term vegetation data instead provide insight into population persistence under interannual variation in climate, land use and biotic conditions (van der Plas *et al*., 2020; de Bello *et al*., 2021; Sperandii *et al*., 2022).

Here, we focus on complementary belowground functions of the root traits. Specific root length (SRL), average root diameter (AD) and root hair incidence (RHI) capture alternative aspects of soil exploration and belowground absorptive strategies (Bergmann *et al*., 2020, 2026). Root tissue density (RTD) reflects conservative tissue construction, whereas root nitrogen concentration (N) reflects acquisitive metabolic investment; both traits are associated with belowground resource economics (Bergmann *et al*., 2020; Freschet *et al*., 2021; Matthus *et al*., 2025). Evaluating these traits separately allows us to test which complementary aspects of belowground function are most strongly associated with long-term population temporal stability.

Land use provides an important environmental context for testing trait–stability relationships in managed semi-natural grasslands. In grasslands where we conducted this research, land-use intensity is quantified as a composite index integrating fertilization, mowing and grazing (Blüthgen *et al*., 2012). These components affect populations through partly different pathways: fertilization mainly alters nutrient availability and productivity, whereas mowing and grazing impose repeated biomass removal, disturbance and changes in vegetation structure (Socher *et al*., 2012; Allan *et al*., 2015). By changing resource limitation, disturbance regimes and competitive conditions, land use may modify how root traits translate into temporal population stability. Testing land use as a moderator therefore allows us to evaluate whether belowground trait-stability relationships shift along management gradients rather than remaining fixed across grasslands (Figure 1).

**Figure 1.**
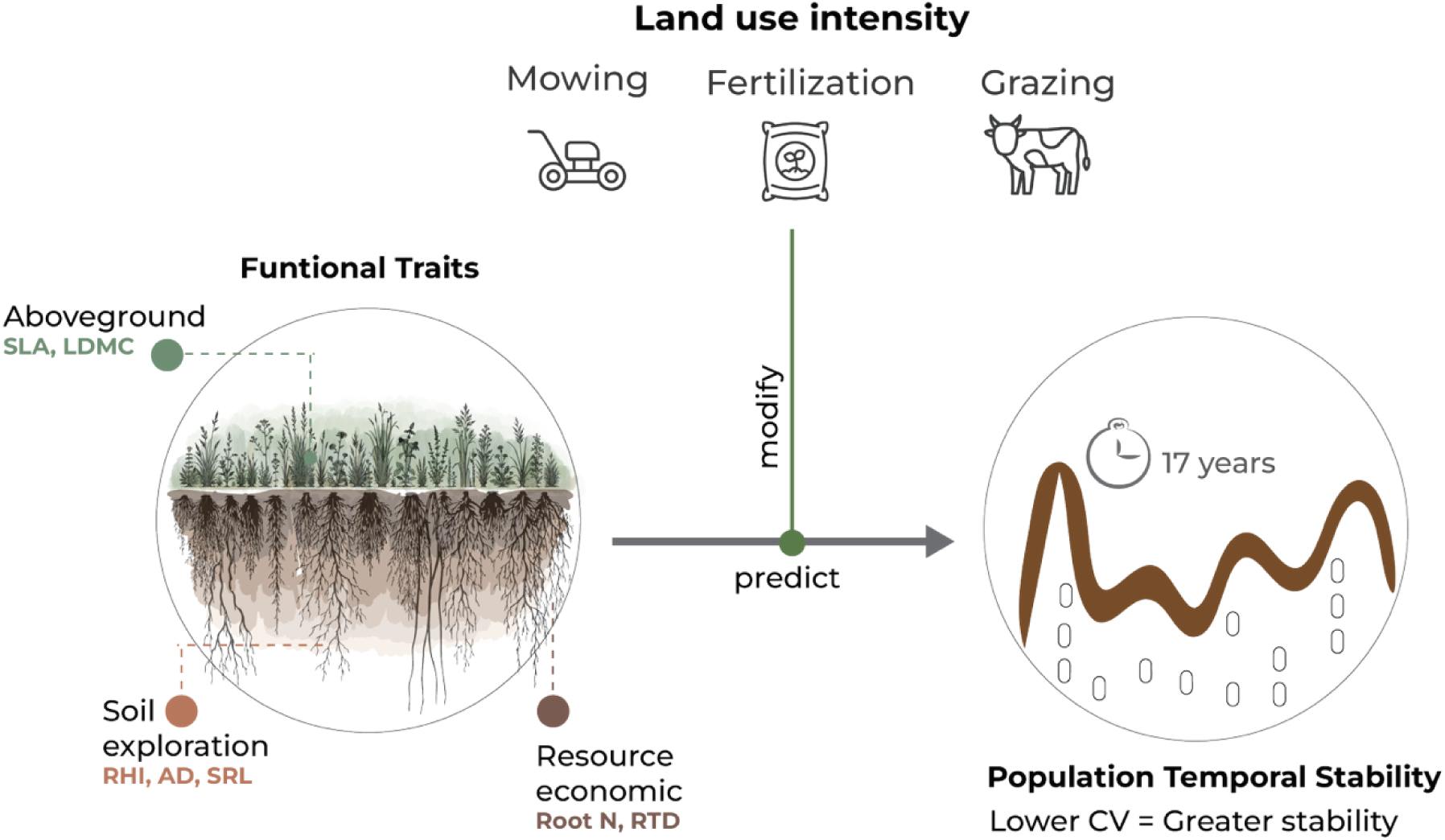
Conceptual overview of the study framework. We tested associations between functional traits and population temporal stability in temperate grasslands and whether these relationships were modified by land use. Traits were grouped into aboveground traits (SLA, specific leaf area; LDMC, leaf dry matter content), soil-exploration traits (RHI, root hair incidence; AD, average root diameter; SRL, specific root length), and root resource-economics traits (root N, root nitrogen concentration; RTD, root tissue density). Land use comprised mowing, fertilization and grazing. Temporal stability was quantified from 17 years of species abundance data, with lower temporal CV indicating higher stability.

Here, we used 17 years of vegetation data from 150 temperate grassland plots, combined with functional trait data for 72 common grassland species, to address three questions.

- First, do root traits predict long-term population temporal stability, and do they explain variation in stability beyond aboveground leaf economics traits?
- Second, which root traits and related functions are the main drivers of population temporal stability?
- Third, does land-use intensity, and its components of fertilization, mowing and grazing, modify the relationship between root traits and temporal stability?

## Materials and Methods

### Study system

The study was conducted as part of the Biodiversity Exploratories, large-scale and long term land-use experiment with 150 grassland plots located in three areas in Germany (www.biodiversity-exploratories.de; Fischer et al. 2010); AEG plots are located in the UNESCO Biosphere Reserve Schwäbische Alb (Swabian Jura) in the southeast of Germany on a limestone-rich mid-mountain range (460–860 m a.s.l.), with Leptosols and Cambisols soil types and a mean annual precipitation (MAP) of 700–1000 mm. HEGs are located in the National Park Hainich-Dün in central Germany on hilly terrain (258–550 m a.s.l.), with Cambisols, Vertisols, and Stagnosols soil types and a MAP of 500–800 mm. SEG plots are located in the UNESCO Biosphere Reserve Schorfheide-Chorin in the northeast of Germany on plain, undulating moraine hills (3–140 m a.s.l.), with Histosols, Luvisols, Gleysols, Cambisols, and Albeluvisols soil types, and a MAP of 480–580 mm.

### Long-term abundance data

To measure temporal variation, we used 17 years (2008–2024) of annual grassland species abundance data from 150 plots (50 per region). In each plot, species cover (%) was recorded during peak biomass (late May–early June) within a permanent 4 × 4 m area (CoreBotany project; Hinderling & Keller, 2025). Temporal stability was estimated as the Coefficient of Variation (CV) in species cover (%) across years, calculated as the standard deviation divided by the mean (Doak *et al*., 1998; Májeková *et al*., 2014; Lepš *et al*., 2018).

### Traits

Functional trait data were obtained from a standardized greenhouse experiment conducted in 2018 on grassland species from the Biodiversity Exploratories (Bergmann *et al*., 2026) In this experiment, species were grown in sterile substrate inoculated with arbuscular mycorrhizal fungi to approximate natural biotic conditions, and root and leaf traits were measured following standardized protocols (Bergmann *et al*., 2026).

Root trait selection was based on complementary belowground functions. We grouped traits into two functional components. Soil exploration; was represented by specific root length (SRL; m g⁻¹), root-hair incidence (RHI; %), and average root diameter (AD; mm), which capture different strategies of soil exploration and root–soil contact (Freschet *et al*., 2021; Matthus *et al*., 2025; Bergmann *et al*., 2026). Root resource economics; was represented by root tissue density (RTD; g cm⁻³) and root nitrogen concentration (N; %). RTD reflects investment in dense, persistent root tissue, whereas root N reflects metabolic investment associated with a more acquisitive resource-use strategy; thus, these traits represent contrasting aspects of the acquisition–conservation trade-off. In addition, we included the aboveground leaf economics traits specific leaf area (SLA; m² kg⁻¹) and leaf dry matter content (LDMC; mg g⁻¹) for comparison with belowground traits (Wright *et al*., 2004; Díaz *et al*., 2016; Conti *et al*., 2023). Multivariate coordination among the seven traits was assessed using principal component analysis (Fig. S1, Fig. S2). All traits were standardized to z-scores prior to analysis.

### Land-use intensity

Land-use intensity was quantified annually using standardized questionnaires completed by farmers and landowners (Blüthgen *et al*., 2012). Three management components were recorded annually: fertilization intensity (kg N ha⁻¹ yr⁻¹), mowing frequency (cuts yr⁻¹), and grazing intensity (livestock units × grazing days ha⁻¹ yr⁻¹). Each component was standardized relative to its regional mean and combined into the dimensionless land-use intensity index (LUI), with higher values indicating more intensive management. The index was square-root transformed following (Blüthgen *et al*., 2012), based on grassland management information from Vogt *et al*., 2019, using the LUI calculation tool of Ostrowski *et al*. 2020. We analyzed both overall LUI and the individual mowing, grazing, and fertilization components to distinguish general from management-specific effects.

### Statistical analyses

To characterize trait coordination, we performed principal component analysis (PCA) on standardized species-level values of the seven aboveground and root traits, using each species once in the analysis (Fig. S1). We additionally calculated pairwise Pearson correlations among the traits to assess the strength and direction of associations between individual traits (Fig. S2).

To test associations between individual functional traits and population temporal stability, we fitted separate linear mixed-effects models for each of the seven traits, with the coefficient of variation in species cover (CV; lower values indicating higher temporal stability) as the response variable (Table S1). Trait, region, and their interaction (*trait × region*) were included as fixed effects, while species identity and plot identity were included as random intercepts to account for repeated observations and spatial structure. Models were fitted using restricted maximum likelihood (REML) in *lme4*, with *lmerTest* used for inference.

To evaluate the relative contribution of correlated traits, we used regularized regression with an L1 penalty (LASSO), an approach suited to datasets with multiple correlated predictors and useful for identifying the relative importance of individual traits (Tibshirani, 1996; Friedman *et al*., 2010). We fitted a single regional model with separate regional intercepts and region-specific coefficients for each of the seven traits. Regional intercepts were unpenalized, while trait coefficients were penalized. The penalty parameter (*λ*) was selected by 10-fold cross-validation, and non-zero coefficients were used to identify traits associated with variation in CV within each region.

To test whether land use modified trait–stability relationships, we fitted separate linear mixed-effects models for each of the seven traits with overall LUI and its grazing, mowing, and fertilization components (Blüthgen *et al*., 2012). For each plot, annual land-use indices were averaged across 2006–2024 and the resulting plot-level means were standardized before analysis. For each *trait × land-use* combination, we compared an additive model containing trait, land use, and region as fixed effects with a model additionally containing the *trait × land-use* interaction. Species and plot identities were included as random intercepts. Models were fitted by maximum likelihood using *lme4* (Bates *et al*., 2014), and interaction significance was assessed using likelihood-ratio tests between the additive and interaction models.

All analyses were conducted in R version 4.3.1 (R Core Team, 2023).

## Results

### Trait coordination

Trait coordination supported a multidimensional root trait coordination rather than a single belowground axis (Figs. S1, S2). PC1 explained 39.7% of total trait variation (λ = 2.78) and PC2 explained 23.4% (λ = 1.64), together accounting for 63.1% of variation; both components had λ > 1 (Fig. S1). PC1 contrasted AD, root N and SLA with RHI, LDMC and SRL, whereas PC2 was characterized mainly by opposing loadings of RTD and SRL. Pairwise correlations were generally weak to moderate, with the strongest correlation between SRL and AD (r = −0.73; Fig. S2). Overall, traits were only partly coordinated, supporting their analysis as individual traits.

### Trait associations with temporal stability

Regularized regression showed that belowground traits had the strongest and most consistent associations with temporal stability across regions (Fig. 2). Since lower CV indicates higher temporal stability, negative coefficients indicate that higher trait values were associated with greater stability, whereas positive coefficients indicate lower stability. In the Schwäbische Alb, average root diameter had the largest negative coefficient (β = −0.23), followed by root-hair incidence (β = −0.19) and specific root length (β = −0.17). In Hainich-Dün, root-hair incidence showed the largest negative coefficient (β = −0.24), followed by average root diameter and specific root length (both β = −0.14). In Schorfheide-Chorin, root tissue density had the largest negative coefficient (β = −0.20), followed by specific root length (β = −0.16), and average root diameter and root-hair incidence (both β = −0.11).

**Figure 2.**
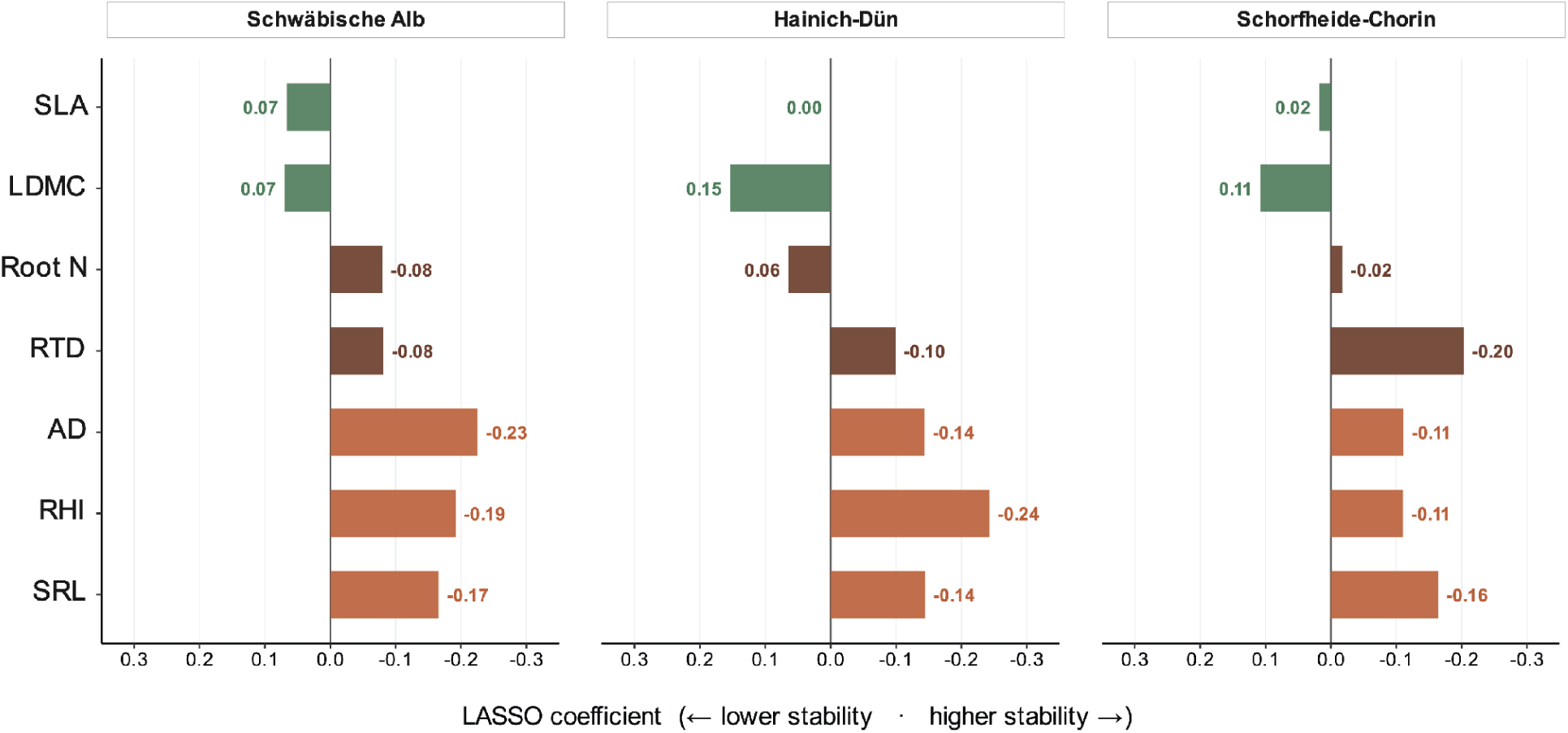
Regional associations of functional traits with population temporal stability. Standardized LASSO coefficients (β) for functional traits predicting species temporal stability (CV) in the Schwäbische Alb (AEG), Hainich-Dün (HEG), and Schorfheide-Chorin (SEG). Traits shown are specific leaf area (SLA), leaf dry matter content (LDMC), root nitrogen content (Root N), root tissue density (RTD), average root diameter (AD), root hair incidence (RHI), and specific root length (SRL). Negative coefficients indicate associations with higher temporal stability (lower CV), whereas positive coefficients indicate associations with lower temporal stability (higher CV). Colors denote functional groups: green = aboveground traits (SLA, LDMC), orange = soil exploration traits (SRL, RHI, AD), and brown = root resource-economics traits (RTD, Root N).

Aboveground traits showed weaker and less consistent associations. Specific leaf area had small positive coefficients in the Schwäbische Alb (β = 0.07) and Schorfheide-Chorin (β = 0.02) and was reduced to zero in Hainich-Dün. Leaf dry matter content showed positive coefficients in all regions, strongest in Hainich-Dün (β = 0.15) and Schorfheide-Chorin (β = 0.11), indicating lower stability with increasing LDMC. Root nitrogen showed small and region-dependent coefficients (β = −0.08, 0.06 and −0.02, respectively).

The single-trait mixed-effects models provided complementary evidence for the importance of individual traits (Table S1). These models evaluated each trait independently, whereas the regularized regression assessed their relative importance while accounting for correlations among traits. Consistent with the regularized regression, SRL, AD and RTD showed stronger associations with temporal stability, whereas RHI and root N showed weaker effects.

### Land-use intensity modulates trait-stability relationships

Land use significantly modified several trait–stability relationships, with the strongest and most widespread shifts occurring along the mowing gradient (Table 1; Fig. 3). Increasing overall LUI, mowing and fertilization strengthened the negative association between root hair incidence (RHI) and CV (β = −0.073, −0.090 and −0.056, respectively; all P ≤ 0.002), indicating that higher RHI became increasingly associated with greater temporal stability under more intensive management. In contrast, the average root diameter (AD) and CV relationship shifted in the positive direction with increasing overall LUI (β = 0.053, P = 0.002), mowing (β = 0.057, P < 0.001) and fertilization (β = 0.034, P = 0.041). Root tissue density (RTD) showed contrasting responses, with its relationship with CV shifting in the positive direction under grazing (β = 0.041, P = 0.033) but becoming more negative under mowing (β = −0.053, P = 0.004). Root N also shifted in the positive direction with overall LUI (β = 0.038, P = 0.023) and mowing (β = 0.049, P = 0.004), while LDMC showed a positive interaction with grazing (β = 0.036, P = 0.033). No significant land-use interactions were detected for SLA or SRL.

**Table 1.** Interaction effects of land-use intensity on trait–stability (CV) relationships. Values are trait × land-use interaction coefficients (β) from linear mixed-effects models with associated P-values; trait and land-use variables were standardized before analysis. Negative coefficients indicate that the trait–CV relationship becomes more negative with increasing land-use intensity, whereas positive coefficients indicate a shift in the positive direction. Bold values indicate significant interactions (P < 0.05).

|  | *OVERALL LUI* |  | *GRAZING* |  | *MOWING* |  | *FERTILISATION* |  |
| --- | --- | --- | --- | --- | --- | --- | --- | --- |
| | $\beta$ | *P-value* | $\beta$ | *P-value* | $\beta$ | *P-value* | $\beta$ | *P-value* |
| **LDMC** | 0.010 | *0.564* | **0.036** | **0.033** | −0.025 | *0.155* | 0.008 | *0.643* |
| **SLA** | 0.001 | *0.951* | −0.014 | *0.407* | 0.015 | *0.389* | 0.003 | *0.860* |
| **RHI** | **−0.073** | **<0.001** | 0.020 | *0.239* | **−0.090** | **<0.001** | **−0.056** | **0.002** |
| **AD** | **0.053** | **0.002** | −0.012 | *0.500* | **0.057** | **<0.001** | **0.034** | **0.041** |
| **SRL** | −0.029 | *0.088* | 0.003 | *0.879* | −0.025 | *0.140* | −0.017 | *0.288* |
| **RTD** | −0.017 | *0.350* | **0.041** | **0.033** | **−0.053** | **0.004** | −0.025 | *0.168* |
| **N** | **0.038** | **0.023** | −0.008 | *0.639* | **0.049** | **0.004** | 0.026 | *0.107* |

**Figure 3.**
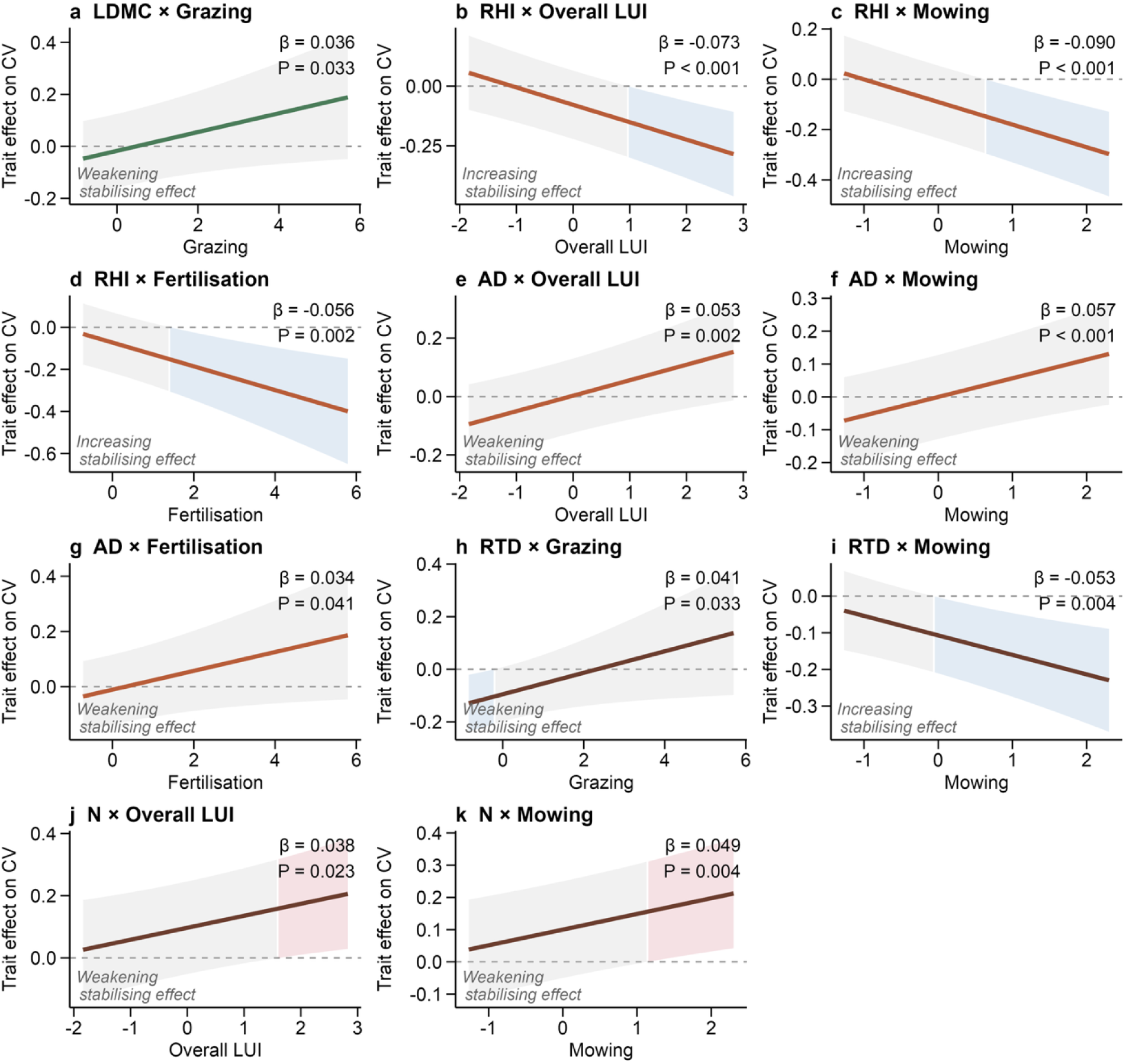
Land-use intensity modulates trait–stability relationships. Conditional trait effects on temporal variability (CV) are shown for significant trait × land-use interactions across overall land-use intensity, grazing, mowing and fertilization gradients. Solid lines show modelled trait effects, dashed lines indicate no effect, and ribbons show 95% confidence intervals. Blue shading indicates significantly negative effects on CV (higher stability), red shading significantly positive effects (lower stability). Interaction coefficients (β) and P-values are shown in each panel.

## Discussion

Our results show that root functional traits are stronger and more consistent predictors of population temporal stability than aboveground leaf economics traits. Across three distinct regions of temperate semi-natural grasslands, the strongest associations with greater population temporal stability were consistently found for root traits. Higher average root diameter, root hair incidence, specific root length and root tissue density were generally associated with higher population temporal stability, although the strength of these relationships varied among regions. Among aboveground traits, LDMC showed a comparatively strong association only in Hainich-Dün, whereas root N was a weak and inconsistent predictor across regions.

### Soil exploration and tissue conservation jointly promote population temporal stability

Soil exploration traits were consistently associated with higher temporal stability across the three grassland regions (AEG, HEG, and SEG). In AEG, stability was most strongly associated with average root diameter, followed by root hair incidence and specific root length, while in HEG, root hair incidence showed the strongest association, followed by average root diameter and specific root length. In SEG, soil exploration traits were also associated with stability, although root tissue density was the strongest predictor. This regional variation suggests that population temporal stability is associated with different architecture of soil exploration rather than with one particular root trait. Greater root diameter, higher root-hair incidence and higher specific root length may represent alternative belowground strategies for maintaining resource access and root–soil interactions under different soil and management conditions. The repeating associations of these traits with stability therefore suggest that the different strategies of soil exploration are important mediators of temporal stability (Weemstra *et al*., 2016; Bergmann *et al*., 2020; Freschet *et al*., 2021)Fig. 2).

Root tissue density represented a more conservative root construction strategy and provided an additional strong association with population temporal stability. Its contribution was particularly pronounced in Schorfheide-Chorin, where it was the strongest predictor of stability. High root tissue density reflects greater investment in dense, persistent root tissues and is often associated with longer root lifespan and slower tissue turnover (Ryser, 1996; Eissenstat & Yanai, 1997). Such investment may help species maintain belowground structures and resource acquisition across years, thereby reducing fluctuations in population abundance. Together with the associations observed for soil-exploration traits, this result suggests that both soil exploration and conservative root construction are relevant to population temporal stability.

Root nitrogen concentration, by contrast, showed only weak and region-dependent associations with population temporal stability. Higher root N is generally associated with the faster, more acquisitive end of the root resource-economics trade-off, whereas high root tissue density represents a more conservative strategy (Freschet *et al*., 2021; Matthus *et al*., 2025; Bergmann *et al*., 2026). In our grasslands, however, root N did not provide a strong stability signal for this acquisitive strategy. This suggests that temporal stability was related more strongly to how roots explore soil and persist through time than to root nitrogen concentration alone.

### Weak contribution of aboveground traits

In contrast, SLA and LDMC, the most commonly used aboveground leaf economics traits in trait-based stability studies (Májeková *et al*., 2014; Bruelheide *et al*., 2018; de Bello *et al*., 2021; Conti *et al*., 2023; Gresse *et al*., 2026; Pan *et al*., 2026), showed weaker and less consistent associations with population temporal stability. Although conservative leaf strategies, characterized by high LDMC or low SLA, are often expected to be associated with higher stability, this relationship does not appear to be universal. LDMC and SLA did not significantly predict ecosystem stability across grassland biodiversity experiments (Craven *et al*., 2018), while recent studies in drylands found that more acquisitive species with higher SLA could show higher population stability (Yan *et al*., 2025; Gresse *et al*., 2026), opposite to the expectation that conservative strategies confer greater stability. In our grasslands, population temporal stability was instead more strongly associated with root traits related to soil exploration and conservative root construction, suggesting that belowground strategies are at least as important as, if not more important than, aboveground strategies in explaining stability.

### Land use shifts mediated root trait-stability relationships

Overall, land use modified which belowground traits mattered most for temporal stability, with the strongest shifts concentrated in soil exploration and root tissue construction traits, particularly along the mowing gradient. Land use did not modify all trait–stability relationships equally. Root hair incidence (RHI), average root diameter (AD), root tissue density (RTD) and root nitrogen concentration (root N) were the root traits whose associations with temporal stability shifted significantly along management gradients, with RHI showing the clearest response. Root hairs increase the root–soil absorptive interface and can enhance access to relatively immobile nutrients (Bates & Lynch, 2000; Haling *et al*., 2013), which may help maintain resource uptake under intensive management. The association of higher RHI with temporal stability strengthened with overall land-use intensity, mowing and fertilization, suggesting that root-hair-mediated soil exploration becomes particularly important where management alters nutrient availability and imposes repeated biomass removal. Average root diameter, another soil-exploration trait, showed an opposite response: its association with temporal stability weakened with increasing overall land-use intensity, mowing and fertilization.

Root tissue density showed management-specific responses, with the association of dense root tissue with stability strengthening under mowing but weakening under grazing. This contrast suggests that different forms of biomass removal may alter the relative importance of conservative root construction, as grazing imposes a more heterogeneous disturbance regime through selective defoliation, trampling and nutrient redistribution (Socher *et al*., 2012; Allan *et al*., 2015). Root N also shifted towards a weaker association with stability under mowing and overall land-use intensity, suggesting that metabolic investment reflected by root N may be less strongly related to stability under more intensive management. Among aboveground traits, only leaf dry matter content (LDMC) showed a significant management-dependent shift, with its association with stability weakening under increasing grazing intensity.

### Toward a belowground framework for temporal stability

Our findings shift the functional interpretation of population temporal stability from a primarily aboveground leaf economic spectrum perspective toward a belowground perspective, where stable populations are defined by the capacity of roots to explore soil space and invest in durable tissues. Our findings identify belowground functional traits as an important component of trait-based studies of population temporal stability.

Here, we focus on well-established fine-root and leaf traits related to resource economics and show that particularly the root traits capture variation in population temporal stability. Future work should combine standardized root trait datasets with targeted field measurements of mycorrhizal and hydrological traits to build a more complete belowground framework for predicting which species remain stable, which decline and how management reshapes grassland population dynamics.

## Conclusion

Overall, this study shows that population temporal stability in grasslands is fundamentally linked to plant belowground strategies. Root traits explain stability beyond what can be inferred from aboveground traits associated with resource economics and reveal multiple stabilizing mechanisms related to soil exploration and conservative root construction. Land use further reshapes these relationships to become more or less stabilizing, especially for root hair incidence, average root diameter, and root tissue density. By placing root traits at the centre of temporal population dynamics, our findings open new research avenues for understanding and predicting population stability in managed grassland ecosystems.

## Acknowledgements

We thank the Plant Ecology group of the University of Tübingen for their valuable input and discussions throughout this project. We thank Lucien D. L. Liancourt for his insightful feedback. We also thank the managers of the three Biodiversity Exploratories, Swabian Alb (Julia Bass, Max Müller), Hainich-Dün (Anna K. Franke), and Schorfheide-Chorin (Franca Marian, Max Müller, Uta Schumacher), and all former managers for their work in maintaining the plot and project infrastructure. We thank Victoria Grießmeier for giving support through the central office, and the respective data manager for managing the central database. We also thank Markus Fischer, Eduard Linsenmair, Dominik Hessenmöller, Daniel Prati, Ingo Schöning, François Buscot, Ernst-Detlef Schulze, Wolfgang W. Weisser, and the late Elisabeth Kalko for their role in setting up the Biodiversity Exploratories project. We thank the administration of the Hainich National Park, the UNESCO Biosphere Reserve Swabian Alb, and the UNESCO Biosphere Reserve Schorfheide-Chorin, as well as all landowners, for their excellent collaboration. Fieldwork permits were issued by the responsible state environmental offices of Baden-Württemberg, Thuringia, and Brandenburg. The work has been funded by the German Research Foundation (DFG) as part of the Priority Programme 1374 “Biodiversity Exploratories” (DFG # 512047507).

## Data availability statement

This work is based on data elaborated by the RootFun project (323522591) and further analyzed within the HAIRphae project (432975993) of the Biodiversity Exploratories program (DFG Priority Program 1374). Vegetation data were obtained from the CoreBotany project (Hinderling & Keller, 2025; Biodiversity Exploratories Information System, Dataset ID 32246). Land-use-intensity indices (LUIs) were calculated as grassland management indicators according to Blüthgen et al. (2012), based on information from landowners on mowing, grazing and fertilization (Vogt et al., 2019), using the LUI calculation tool implemented in BExIS (Ostrowski et al., 2020). All data were generated within the Biodiversity Exploratories programme.

## Supplementary Material

**Table S1.** Linear mixed-effects model results for each trait predicting species temporal stability (CV of population abundances) across grassland regions. Each trait was modelled separately with region as a fixed moderator and species identity and plot as random intercepts (reference region = AEG). All traits were standardized prior to analysis. Lower CV indicates higher temporal stability. Bold values indicate P < 0.05.

| TRAIT | REGION | B | SE | T | P-VALUE |
| --- | --- | --- | --- | --- | --- |
| **SLA** (*Aboveground*) | AEG | -0.037 | 0.061 | -0.60 | 0.548 |
|  | HEG | 0.022 | 0.040 | 0.56 | 0.575 |
|  | SEG | -0.019 | 0.048 | -0.41 | 0.682 |
| **LDMC** (*Aboveground*) | AEG | -0.009 | 0.078 | -0.11 | 0.912 |
|  | HEG | -0.048 | 0.040 | -1.18 | 0.237 |
|  | SEG | 0.050 | 0.045 | 1.12 | 0.263 |
| **SRL** (*Soil exploration*) | AEG | **0.193** | 0.070 | 2.77 | **0.007** |
|  | HEG | **-0.115** | 0.041 | -2.82 | **0.005** |
|  | SEG | **-0.133** | 0.046 | -2.91 | **0.004** |
| **RHI** (*Soil exploration*) | AEG | -0.067 | 0.080 | -0.84 | 0.402 |
|  | HEG | -0.039 | 0.042 | -0.94 | 0.349 |
|  | SEG | 0.062 | 0.049 | 1.27 | 0.203 |
| **AD** (*Soil exploration*) | AEG | -0.065 | 0.071 | -0.91 | 0.364 |
|  | HEG | 0.055 | 0.039 | 1.41 | 0.157 |
|  | SEG | **0.111** | 0.048 | 2.30 | **0.021** |
| **RTD** (*Resource-economics*) | AEG | **-0.116** | 0.059 | -1.98 | **0.050** |
|  | HEG | 0.033 | 0.040 | 0.82 | 0.410 |
|  | SEG | 0.042 | 0.051 | 0.82 | 0.413 |
| **N** (*Resource-economics*) | AEG | 0.042 | 0.081 | 0.51 | 0.610 |
|  | HEG | **0.081** | 0.039 | 2.08 | **0.038** |
|  | SEG | 0.068 | 0.045 | 1.50 | 0.133 |

**Figure S1.**
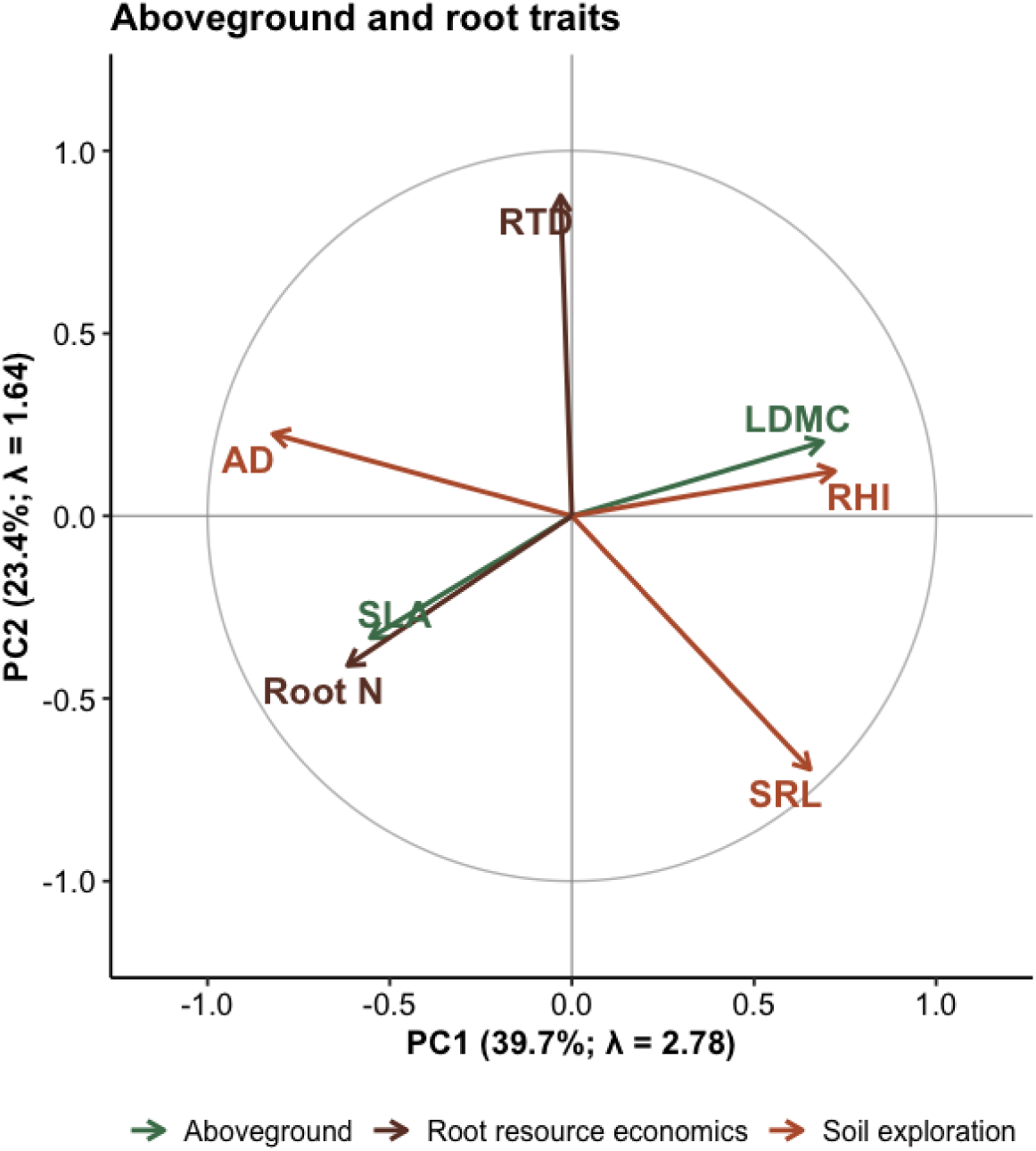
Principal component analysis (PCA) of aboveground and root functional traits. Arrows represent correlations of individual traits with the first two principal components; arrow direction indicates the sign of the association and arrow length its strength. Traits are grouped as aboveground (SLA, LDMC), soil exploration traits (SRL, RHI, AD), and root resource-economics traits (RTD, root N). PC1 explained 39.7% of total trait variation (λ = 2.78) and PC2 explained 23.4% (λ = 1.64), together accounting for 63.1% of total variation. Both components had eigenvalues > 1. Colors denote functional groups: green = aboveground traits (SLA, LDMC), orange = soil exploration traits (SRL, RHI, AD), and brown = root resource-economics traits (RTD, Root N).

**Figure S2.**
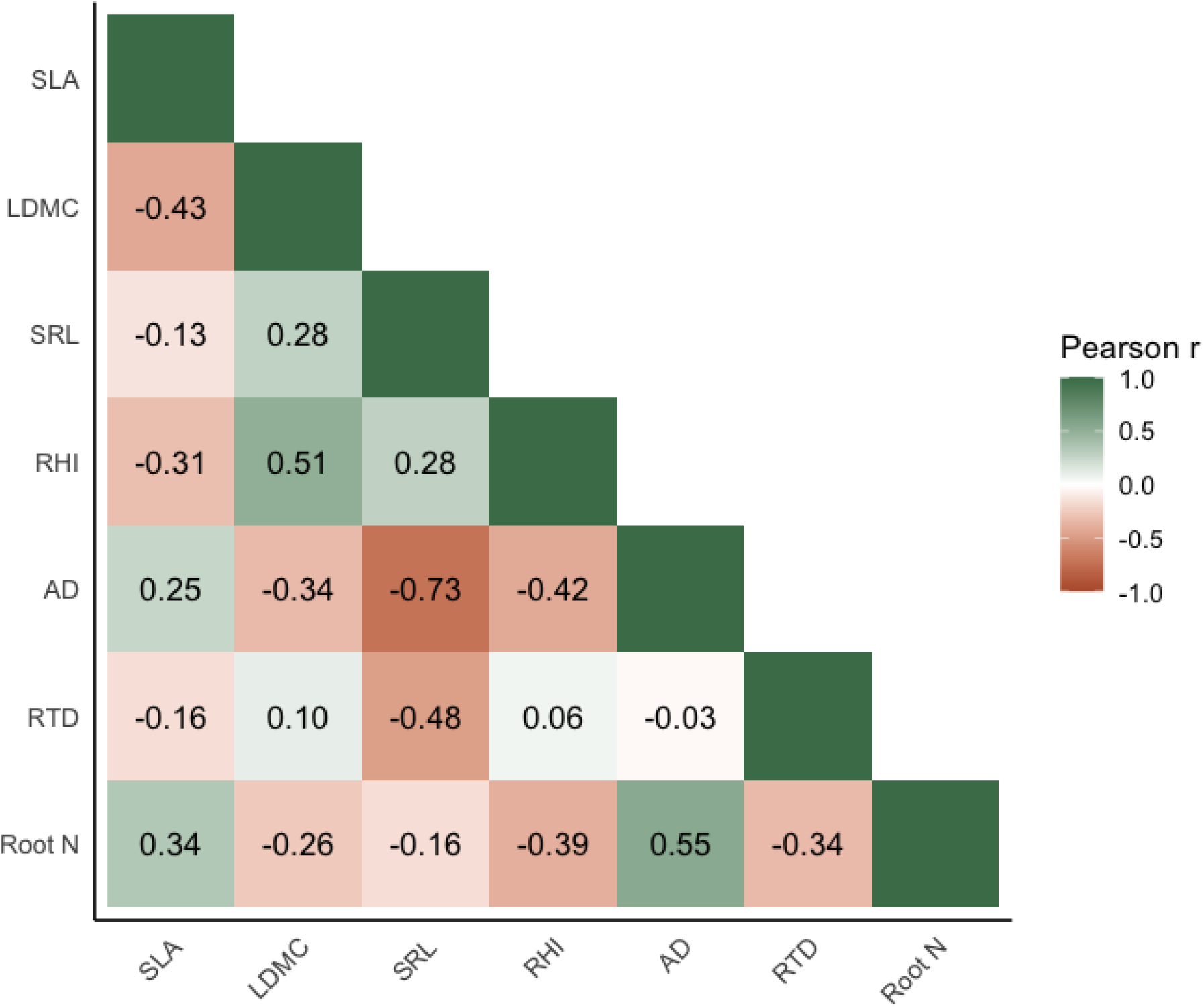
Correlation matrix of belowground and aboveground functional traits. Values indicate pairwise Pearson correlation coefficients (r) among the seven traits. Green indicates positive correlations and orange indicates negative correlations; darker shading represents stronger correlations.

**Table S2.** The three study regions within the Biodiversity Exploratories. Environmental and geomorphological characteristics of the three regions. (Fischer et al. 2010).

|  | *Schwäbische Alb*<br>*(AEG)* | *Hainich-Dün*<br>*(HEG)* | *Schorfheide-Chorin*<br>*(SEG)* |
| --- | --- | --- | --- |
| ***Annual precipitation***<br>***(mm)*** | 700–1000 | 500–800 | 480 (East) – 580 (West) |
| ***Elevation (m a.s.l.)*** | 460–860 | 285–550 | 3–140 |
| ***Area (km<sup>2</sup>)*** | ~422 | ~130 | ~1300 |
| ***Dominant soil types*** | Leptosols (steep slopes), Cambisols | Eutric Cambisols, Stagnosols (on loess over Triassic limestone), Vertisols | Histosols (drained), Luvisols, Gleysols, Cambisols, Albeluvisols |
| ***Geological bedrock*** | Loess over calcareous Jurassic shell limestone; grasslands rich in clay | Calcareous bedrock with karst features | Glacial till, often covered by glaciofluvial or aeolian sands; fertile loamy soils in grasslands |
| ***Geomorphology*** | Mid-mountain range | Hilly terrain (Central German hilly lands) | Plain to gently undulating moraine landscape with fens and wetlands in depressions |

